# Plasma concentrations of anti-inflammatory cytokine TGF-β are associated with hippocampal structure related to explicit memory performance in older adults

**DOI:** 10.1101/2022.12.21.521442

**Authors:** Matthias Raschick, Anni Richter, Larissa Fischer, Lea Knopf, Annika Schult, Renat Yakupov, Gusalija Behnisch, Karina Guttek, Emrah Düzel, Ildiko Rita Dunay, Constanze I. Seidenbecher, Burkhart Schraven, Dirk Reinhold, Björn H. Schott

**Author notes:** These authors contributed equally to this work (Matthias Raschick, Anni Richter). These authors share senior authorship (Dirk Reinhold, Björn H. Schott). Address for correspondence: PD Dr. Dr. Björn Hendrik Schott, Leibniz Institute for Neurobiology, Brenneckestr. 6, 39118 Magdeburg, Germany, /.

## Abstract

Human cognitive abilities, and particularly hippocampus-dependent memory performance typically decline with increasing age. Immunosenescence, the age-related disintegration of the immune system, is increasingly coming into the focus of research as a considerable factor contributing to cognitive decline. In the present study, we investigated potential associations between plasma levels of pro- and anti-inflammatory cytokines and learning and memory performance as well as hippocampal anatomy in young and older adults. Plasma concentrations of the inflammation marker CRP as well as the pro-inflammatory cytokines IL-6 and TNF-α and the anti-inflammatory cytokine TGF-β_1_ were measured in 142 healthy adults (57 young, 24.47 ± 4.48 years; 85 older, 63.66 ± 7.32 years) who performed tests of explicit memory (Verbal Learning and Memory Test, VLMT; Wechsler Memory Scale, Logical Memory, WMS) with an additional delayed recall test after 24 hours. Hippocampal volumetry and hippocampal subfield segmentation were performed using FreeSurfer, based on T1-weighted and high-resolution T2-weighted MR images. When investigating the relationship between memory performance, hippocampal structure, and plasma cytokine levels, we found that TGF- β_1_ concentrations were positively correlated with the volumes of the hippocampal CA4-dentate gyrus region in older adults. These volumes were in turn positively associated with better performance in the WMS, particularly in the delayed memory test. Our results support the notion that endogenous anti-inflammatory mechanisms may act as protective factors in neurocognitive aging.

## 1. Introduction

Structural and functional alterations of the hippocampus-dependent memory system in aging are a well-replicated finding (Nyberg 2017; Gorbach et al. 2017), and there is mounting evidence for age-related immune dysregulation as a potential factor contributing to age-related cognitive decline and ultimately clinically relevant memory disorders (Brosseron et al. 2022). Normal aging, while not usually considered a disease (Rattan 2014), is accompanied by a low-grade pro-inflammatory state commonly termed “inflammaging” (Franceschi et al. 2000). Lifelong exposure to an accumulating antigen load is thought to act as a chronic stressor on the immune system and to promote chronic, asymptomatic, inflammatory activity (De Martinis et al. 2005). Aging cells frequently enter a state of senescence, which is characterized by a silenced cell cycle and a proinflammatory phenotype (“senescence-associated secretory phenotype”, SASP). SASP is defined by senescence-induced increased secretion of pro-inflammatory molecules like cytokines or chemokines (Basisty et al. 2020) (see http://www.saspatlas.com/), which in turn induce senescence in neighboring cells (Nelson et al. 2012; Acosta et al. 2013). Furthermore, reduced adaptive immunity in old age is thought to further stimulate activity of the innate immune system as a compensatory mechanism (Fülöp et al. 2017). The resulting pro-inflammatory state increases the risk for age-related diseases like cancer or cardiovascular disease and for cognitive dysfunction (Tracy 2003; Akbaraly et al. 2013). Notably, some individuals show relatively preserved memory performance in old age (Nyberg and Pudas 2019), and research in over-centenarians suggests that balanced anti-inflammatory activity in addition to the age-typical pro-inflammatory phenotype may constitute a protective factor counteracting the negative effects of inflammaging (Franceschi 2007).

A prominent anti-inflammatory cytokine is the transforming growth factor β (TGF-β). Mammalian TGF-β protein exists in three different isoforms (TGF-β_1_, TGF-β_2_ and TGF-β_3_), with TGF-β_1_ being most abundant (Dobaczewski et al. 2011). TGF-β_1_ exerts considerable antiinflammatory and neuroprotective effects (Preller et al. 2007; Harder et al. 2014), and there is evidence that TGF-β_1_ is further involved in plasticity processes related to learning and memory. For example, mice treated with TGF-β_1_ before being exposed to amyloid-β exhibit less synaptic loss and better-preserved memory (Diniz et al. 2017), and, in rats, TGF-β_1_ protects against amyloid-β-dependent neurodegeneration and release of pro-inflammatory cytokines (Shen et al. 2014; Chen et al. 2015). Additionally, TGF-β_1_ can induce the regrowth of damaged neurons after axonal injury (Abe et al. 1996). TGF-β_1_ knock-out mice express lower levels of the synaptic marker synaptophysin and the plasticity-related protein laminin as well as lower synaptic density in hippocampal neurons (Brionne et al. 2003). Inhibition of TGF-β_1_ signaling pathways in the mouse hippocampus leads to reduced expression of long-term potentiation (LTP) and impaired object recognition (Caraci et al. 2015), and further studies suggest that TGF-β_1_ is involved in the expression of the late phase of LTP (Mikheeva et al. 2019; Nenov et al. 2019; Caraci et al. 2015). Despite considerable evidence for a contribution of TGF-β_1_ to cellular mechanisms of learning and memory in rodents, little is known about a potential role of TGF-β_1_ in human neural plasticity and memory. There is, however, evidence for a protective role of TGF-β_1_ in neurodegenerative and neuroinflammatory diseases, particularly in Alzheimer’s disease and multiple sclerosis (Diniz et al. 2017; Martínez-Canabal 2015).

Based on the well-documented role of TGF-β_1_ in rodent hippocampal plasticity and memory and its neuroprotective effects in human disease, we aimed to assess a potential relationship between TGF-β_1_ concentrations, hippocampal structure and learning and memory performance in healthy humans, with a focus on older adults. To control for concurrent pro-inflammatory activity, we additionally determined plasma levels of the inflammation marker C-reactive protein (CRP) and of the pro-inflammatory cytokines interleukin 6 (IL-6) and tumor necrosis factor a (TNF-a). As cognitive measures, we employed well-established neuropsychological tests of explicit memory (Verbal Learning and Memory Test, VLMT; Wechsler Memory Scale, WMS) and performed automated hippocampal segmentation and volumetry of hippocampal subfields (Quattrini et al. 2020) using magnetic resonance imaging (MRI) in a previously described cohort of young and older adults (Soch et al. 2021b; Soch et al. 2021a). We hypothesized that higher TGF-β_1_ plasma concentrations would be associated with larger hippocampal volumes and better memory performance, particularly in older adults. Since hippocampal neurogenesis occurs primarily in the subgranular zone (SGZ) of the dentate gyrus (DG) and the functionally related CA4 region (Bond et al. 2021), we additionally conducted a focused analysis in those subregions. According to previous studies showing that the volumes of the input regions of the hippocampus (i.e., DG, CA4 and CA3 regions) are positively correlated with verbal memory (Mueller et al. 2011; Aslaksen et al. 2018; Travis et al. 2014; Kirchner et al. 2023), we further hypothesized that larger volumes of the CA4 region and DG would correlate with better memory performance in the VLMT and WMS.

## 2. Materials and Methods

### 2.1 Study Cohort

#### 2.1.1 Recruitment

Participants were recruited via local press in Magdeburg, the online presence (www.lin-magdeburg.de) and social media of the Leibniz Institute for Neurobiology, as well as flyers in shopping centers and at Otto von Guericke University events. Individuals interested in participating provided information about potential contraindications for participation via telephone interview before they were invited to the study. All procedures were approved by the Ethics Committee of the Medical Faculty of the Otto von Guericke University Magdeburg and conducted according to their guidelines (ethics approval number 33/15). All subjects provided written informed consent to participate in accordance with the Declaration of Helsinki (World Medical Association Declaration of Helsinki: ethical principles for medical research involving human subjects, 2013).

#### 2.1.2 Participants

In the present study, we investigated participants from a previously described cohort consisting of neurologically and psychiatrically healthy young and older adults (Table 1) who underwent multimodal phenotyping, including neuropsychology, MRI, and investigation of blood-based biomarkers as part of the *Autonomy in Old Age* research alliance. A detailed characterization of the cohort and description of testing procedures has been reported previously (Soch et al. 2021b; Soch et al. 2021a; Richter et al. 2023). All participants were right-handed according to selfreport. The Mini-International Neuropsychiatric Interview (M.I.N.I.; Sheehan et al. 1998; German version by Ackenheil et al. 1999) was used to exclude present or past psychiatric disorders. Further contraindications for participation included alcohol or drug abuse, the use of neurological or psychiatric medication, and major psychosis (schizophrenia, bipolar disorder) in a first-degree relative. For the purpose of the present study, additional contraindications were chronic infectious, autoimmune or other inflammatory diseases (e.g., Crohn’s disease, rheumatic diseases, Hashimoto’s thyroiditis, or celiac disease) as well as regular use of immunomodulatory or anti-inflammatory medication. After excluding participants with missing data, a total of 142 participants (57 young, 85 older) were available for data analysis.

**Table 1:** Demographics, cytokine plasma concentrations, hippocampal volumes, and memory performance

| | older ($\geq 50$ yrs) | young (18-35 yrs) | Age effects |
| --- | --- | --- | --- |
| Size of cohort | n = 85 | n = 57 |  |
| Age distribution | 63.66 ± 7.32 | 24.47 ± 4.48 | |
| Sex distribution<br>(f / m) | 53 / 32 | 34 / 23 | $\chi^2 = 0.11$;<br>p = .740 |
| IL-6 [pg/ml] | **1.50 ± 0.99** | **0.85 ± 0.64** | $t_{139.8} = 4.71$;<br>p < .001 |
| TNF-α [pg/ml] | **0.39 ± 0.15** | **0.27 ± 0.21** | $t_{94.5} = 3.70$;<br>p < .001 |
| TGF-$\beta_1$ [pg/ml] | **2897.37 ± 1517.86** | **3300.14 ± 1806.62** | $t_{140} = 1.44$;<br>p = .154 |
| CRP [ng/ml] | **2622.00 ± 2202.47** | **2416.8 ± 3059.14** | $t_{140} = .47$;<br>p = .643 |
| **whole HC** |  |  |  |
| left | **3210.00 ± 292.16** | **3400.27 ± 293.84** | $t_{140} = 3.80$<br>p < .001 |
| right | **3298.54 ± 316.87** | **3452.24 ± 303.19** | $t_{140} = 2.88$<br>p = .005 |
| **DG** |  |  |  |
| left | **276.89 ± 32.19** | **300.27 ± 35.13** | $t_{140} = 4.09$<br>p < .001 |
| right | **289.82 ± 36.34** | **302.53 ± 36.89** | $t_{140} = 2.03$<br>p = .044 |
| **CA4** |  |  |  |
| left | **236.20 ± 25.17** | **248.24 ± 26.89** | $t_{140} = 2.72$<br>p = .007 |
| right | **247.09 ± 28.71** | **250.04 ± 30.25** | $t_{140} = .59$<br>p = .558 |
| **VLMT** |  |  |  |
| learning | **52.35 ± 9.24** | **68.23 ± 4.78** | $t_{132.7} = 13.39$<br>p < .001 |
| 24h recall | **8.89 ± 3.53** | 14.26 ± 1.23 | $t_{111.7} = 12.89$<br>p < .001 |
| **WMS** |  |  |  |
| immediate recall | **25.58 ± 6.54** | **32.30 ± 6.98** | $t_{140} = 5.84$<br>p < .001 |
| 24h recall | **21.74 ± 7.18** | **30.14 ± 7.81** | $t_{140} = 6.60$<br>p < .001 |

Sex (female/male) and age as well as cytokine and CRP concentrations (mean ± standard deviation) are shown. HC: hippocampus; DG: dentate gyrus; VLMT: Verbal Learning and Memory Test; WMS: Wechsler Memory Scale (Logical Memory subscale).

### 2.2. Cognitive testing

#### 2.2.1 General procedure of testing

After signing consent and privacy forms, participants completed questionnaires on general health and MRI contraindications, followed by the M.I.N.I. (see above). Participants older than 50 years additionally underwent the Mini Mental State Examination (MMSE) (Folstein et al. 1975) to rule out MCI or dementia. Afterwards, all participants completed the Multivocabulary Intelligence Test (MWT-B) (Lehrl 2005). This was followed by the actual computer-based cognitive test battery, which included a German version of the VLMT (see 2.2.2) (Helmstaedter 2001) and a German auditory version of the WMS for logical memory (see 2.2.3) (Härting et al. 2000). For a comprehensive description of the test battery see (Richter et al. 2023).

#### 2.2.2 Verbal Learning and Memory Test (VLMT)

The VLMT includes two lists of 15 semantically unrelated words, a study list and a distracter list (Helmstaedter 2001). The experiment was divided into a learning phase and a recall phase. In the learning phase, the words of the first list were presented consecutively visually. After the presentation of all words, participants were asked to write down each word they could remember. This procedure was repeated five times in succession. Next, a second list was presented once, followed by a written recall. This list served as a distractor list and was followed by the recall phase, in which the words from the first list were to be written down. Further recall phases followed after 30 minutes and after 24 hours.

#### 2.2.3 Wechsler Memory Scale (WMS): Logical Memory

The subscale *Logical Memory* of the WMS was implemented as a slightly modified, auditory version (Härting et al. 2000). The test persons listened to two short stories over headphones, which they were asked to write down immediately after listening. Recall tests took place 30 minutes and 24 hours later. The recalled stories were rated by two independent experimenters according to an evaluation sheet with 25 items (i.e., details from the stories), and thus a maximum of 25 points could be obtained per story and recall delay.

### 2.3 Determination of TGF-β plasma levels

#### 2.3.1 Sample collection, processing and storage

Venous blood samples were collected from all participants by healthcare professionals after informed consent, including two plasma tubes with sodium citrate as anticoagulant. Platelet-poor citrate plasma was prepared using a standardized two-step separation method (Reinhold et al. 1997). Citrate plasma samples were stored as aliquots in Eppendorf tubes at −80 °C until measurement. Whenever the interval between the two test days was seven days or more, a blood sample was taken for both test days.

#### 2.3.2 Enzyme-linked immunosorbent assay (ELISA)

Plasma concentrations of TGF-β_1_ were determined via enzyme-linked immunosorbent assay (ELISA). Additional ELISA measurements were carried out to determine plasma levels of CRP and the pro-inflammatory cytokines IL-6 and TNF-α. Commercially available Quantikine^®^ ELISA kits (R&D Systems Inc., Minneapolis, MN, USA) were used for this purpose. For IL-6 and TNF-α, the high-sensitivity variant was used, as low plasma concentrations of these cytokines were expected in healthy individuals, and the high-sensitivity variant is characterized by a lower minimum detection dose.

### 2.4. Magnetic resonance imaging

#### 2.4.1. MRI data acquisition

MRI data were collected using two MRI systems (3 Tesla Verio-Syngo MR system, Siemens Medical Systems, Erlangen, Germany and 3 Tesla Skyra Fit MR system). For volumetric analyses, a T1-weighted 3D Magnetization Prepared Rapid Acquisition Gradient Echo (MPRAGE) image was acquired (TR = 2.5 s, TE = 4.37 ms, flip angle = 7°, 192 sagittal slices, in-plane resolution = 256 x 256, isotropic voxel size = 1 mm^3^). In addition, high-resolution coronal T2-weighted images were acquired using a protocol optimized for medial temporal lobe volumetric analyses (TR = 3.5 s, TE = 353 ms, 64 coronal slices orthogonal to the hippocampal axis, in-plane resolution = 384 x 384, voxel size = 0.5 x 0.5 x 1.5mm^3^) (Richter et al. 2023).

#### 2.4.2. Volumetric measurement of hippocampal subfields

Automated volumetric analysis of the individual participants’ hippocampi and their subfields was performed using FreeSurfer 6.0 (Fischl 2012) and the module for segmentation of hippocampal subfields, which is based on both *in* vivo scans of human participants and *ex vivo* scans of hippocampi from autopsy specimens (Iglesias et al. 2015). Previous analyses have confirmed the robustness of this protocol across different MRI scanners (Quattrini et al. 2020). In addition to the T1-weighted MPRAGE images, high-resolution coronal T2-weighted images (see 2.4.1) were used to improve segmentation accuracy (Dounavi et al. 2020).

### 2.5. Statistical analysis

Statistical analysis was performed using Matlab R2018b (Mathworks, Natick, MA), SPSS Statistics v23 (IBM, Armonk, NY), and R, version 4.0.4 (https://www.r-project.org/), with RStudio version 1.4.1103 (RStudio Team, 2021), employing the R packages psych (Revelle 2022) and sjPlot (Lüdecke 2022).

Main effects of age group and gender on cytokine concentrations, hippocampal volumes, and memory performance were assessed using MANOVAs followed by *post hoc* two-sample t-tests. Whenever Levene’s test was significant, t-tests for unequal variances (i.e., Welch’s tests) were employed.

To investigate a potential direct association of pro- and anti-inflammatory cytokines with learning and memory performance in the VLMT and in the WMS Logical Memory subscale, multiple linear regressions were computed with the concentrations of TGF-β_1_, CRP, IL-6, and TNF-α as well as age and gender as independent variables. An additional independent variable was the infection history of the subjects (i.e., self-report of “cold, flu, urinary tract infection or similar” or vaccination) within four weeks prior to testing (hence referred to as “immune event history”). The dependent variables were learning and memory performance and delayed recall in the VLMT and the WMS. The measure of learning performance for the VLMT was the sum of remembered items from learning sessions 1-5. For the WMS, the sum of the remembered items from both stories in the learning session served as measure of learning performance. Delayed recall performance was quantified as the number of recalled words or story items in the 24-hour recall of the VLMT and WMS, respectively.

Statistical analysis of the relationship between TGF-β plasma concentrations and MRI data (i.e., hippocampal volumes and individual hippocampal subfields) was performed analogously to the behavioral data analysis. Multiple linear regressions were calculated with the concentrations of TGF-β, IL-6, TNF-α, and CRP as well as age, sex, and immune event history as independent variables and the volumes of the hippocampus or hippocampal subregions (DG, CA) as dependent variables, separately for each hemisphere. A correction for the false discovery rate (FDR) was applied for the number of regression analyses (N = 6). To assess for a potential association between hippocampal structure and memory performance, Pearson’s correlations were computed between the volumes of subregions that showed a robust association with cytokine plasma concentrations (p < .05, FDR-corrected) and performance in the VLMT (learning, delayed recall) and in the WMS (immediate and delayed recall).

## 3. Results

### 3.1. Age and gender differences in cytokine levels, hippocampal volumes, and memory performance

Table 1 displays the demographic data, cytokine levels, hippocampal volumes, and performance in the VLMT and WMS, separately for the two age groups. A MANOVA with age and gender as fixed factors and TGF-β1, IL-6, TNF-a and CRP as dependent variables revealed significant effects of age group (Wilks’ *Λ* = .789; F_4,135_ = 9.00; p < .001) and gender (Wilks’ *Λ* = .919; F_4,135_ = 2.97; p = .022) as well as an age by gender interaction (Wilks’ *Λ* = .901; F_4,135_ = 3.67; p = .007). *Post hoc* two-sample t-tests revealed significantly higher levels of IL-6, and TNF-a in older compared to young adults, but no age differences in TGF-β1, or CRP plasma concentrations (Table 1). Significant gender-related differences were observed for CRP in the younger cohort only, with higher levels in women (young: t_37.6_ = 4.17; p < .001; older: t_83_ = .46; p = .636). No gender differences in either age group were found for TGF-β1, IL-6, or TNF-a plasma concentrations (all p > .061).

Volumes of the hippocampus and of CA4 and DG subregions showed highly significant differences as a function of both age group (Wilks’ *Λ* = .618; F_6,133_ = 13.70; p < .001) and gender (Wilks’ *Λ* = .880; F_6,133_ = 3.04; p = .008) as well as a significant age by gender interaction (Wilks’ *Λ* = .905; F_6,133_ = 2.33; p = .036). Table 1 displays the results of the *post hoc* t-tests, which showed that older adults had significantly lower volumes of the whole hippocampus bilaterally as well as of all subregions tested except for the right CA4 region. Regarding gender effects, *post hoc* t-tests in older adults revealed significantly larger volumes of the whole hippocampus and of the subfields in men compared to women (all p < .043). In young adults, the same tendency was observed, but largely only at a trend-level (.019 < p (one-tailed) < .056). This is in line with previous studies showing larger hippocampal volumes in males, which can be attributed to larger intracranial or total brain volumes (Tan et al. 2016).

As expected, memory performance was lower in older compared to young adults. A MANOVA with VLMT learning performance and 24h delayed recall performance as dependent variables revealed a highly significant effect of age group (Wilks’ *Λ* = .459; F_2,137_ = 80.74; p < .001) but no effect of gender and no interaction (all p > .120). *Post hoc* two-sample t-tests showed significantly higher performance in young compared to older adults in both learning and 24h delayed recall (Table 1). Similarly, WMS logical memory performance was significantly lower in older compared to young adults (main effect of age group: Wilks’ *Λ* = .774; F_2,137_ = 20.02; p < .001; see Table 1 for *post hoc* two-sample t-tests), but no effect of gender and no interaction (all p > .307).

### 3.2. Association of TGF-ß plasma concentrations with hippocampal structure

As TGF-β has previously been described to affect hippocampal neurogenesis, which occurs primarily in the SGZ, we hypothesized the subregions of the hippocampus most likely to be related to TGF-β_1_ plasma concentrations would be the DG and the CA4 region. To investigate potential associations between plasma concentrations of TGF-β_1_ or other cytokines (IL-6, TNF-α) and hippocampal structures, we computed multiple linear regressions with the volumes of the entire hippocampus and the DG and CA4 regions as the dependent variable, separately for each hemisphere. Immune event history, age, and gender were included as additional independent variables.

TGF-β_1_ plasma concentrations were significantly positively correlated with the volumes of the CA4 and DG subregions as well as the volumes of the entire hippocampus in both hemispheres (all p < .050, FDR-corrected) in older adults. Other factors influencing the volumes of these regions in older adults were age and gender (all p < .032). Table 2 displays the adjusted regression coefficients and significance levels for all immune markers and anatomical regions, and Figure 1 depicts the correlations between TGF-β_1_ plasma concentrations and hippocampal volumes. However, no effects of TGF-β1 plasma concentrations, age or gender could be observed in young adults (all p > .060, uncorrected), suggesting that the positive relationship between TGF-β_1_ plasma levels and the hippocampal structure was restricted to older adults.

**Table 2:** Immune markers and hippocampal subfield volumes

|  | whole HC |  | CA4 |  | DG |  |
| --- | --- | --- | --- | --- | --- | --- |
|  | left | right | left | right | left | right |
| **TGF-β** |  |  |  |  |  |  |
| std. $\beta$ | **0.43** | **0.22** | **0.32** | **0.32** | **0.27** | **0.29** |
| CI | **[0.11 0.76]** | **[0.03 0.40]** | **[0.13 0.51]** | **[0.13 0.50]** | **[0.07 0.46]** | **[0.10 0.48]** |
| p | **.025\*\*** | **.023\*\*** | **.001\*\*** | **.001\*\*** | **.008\*\*** | **.003\*\*** |
| **IL-6** |  |  |  |  |  |  |
| std. $\beta$ | 0.06 | 0.08 | 0.13 | 0.18 | 0.16 | 0.18 |
| CI | [-0.16 0.29] | [-0.13 0.29] | [-0.09 - 0.35] | [-0.04 0.40] | [-0.06 0.39] | [-0.04 0.40] |
| p | .566 | .465 | .242 | .099 | .154 | .105 |
| **TNF-α** |  |  |  |  |  |  |
| std. $\beta$ | 0.00 | -0.11 | -0.00 | -0.08 | -0.02 | -0.07 |
| CI | [-0.21 0.21] | [-0.32 0.09] | [-0.21 0.21] | [-0.28 0.13] | [-0.24 0.19] | [-0.28 0.14] |
| p | .999 | .285 | .994 | .466 | .833 | .505 |
| **CRP** |  |  |  |  |  |  |
| std. $\beta$ | 0.21 | **0.22** | 0.07 | -0.02 | 0.06 | -0.03 |
| CI | [-0.01 0.43] | **[0.01 0.44]** | [-0.15 0.29] | [-0.24 0.19] | [-0.17 0.28] | [-0.25 0.19] |
| p | .066 | **.039\*** | .504 | .843 | .340 | .793 |
| **immune event** |  |  |  |  |  |  |
| std. $\beta$ | -0.47 | -0.47 | -0.36 | **-0.56** | -0.26 | -0.47 |
| CI | [-0.99 0.06] | [-0.97 0.03] | [-0.88 0.15] | **[-1.07 -0.06]** | [-0.78 0.27] | [-0.98 0.04] |
| p | .080 | .063 | .160 | **.029\*** | .340 | .072 |

Results of multiple regression analyses are shown. std. $\beta$ : standardized regression coefficients. CI: confidence interval. \* $p < .05$ , uncorrected; \*\* $p < .05$ , FDR-corrected.

**Figure 1:**
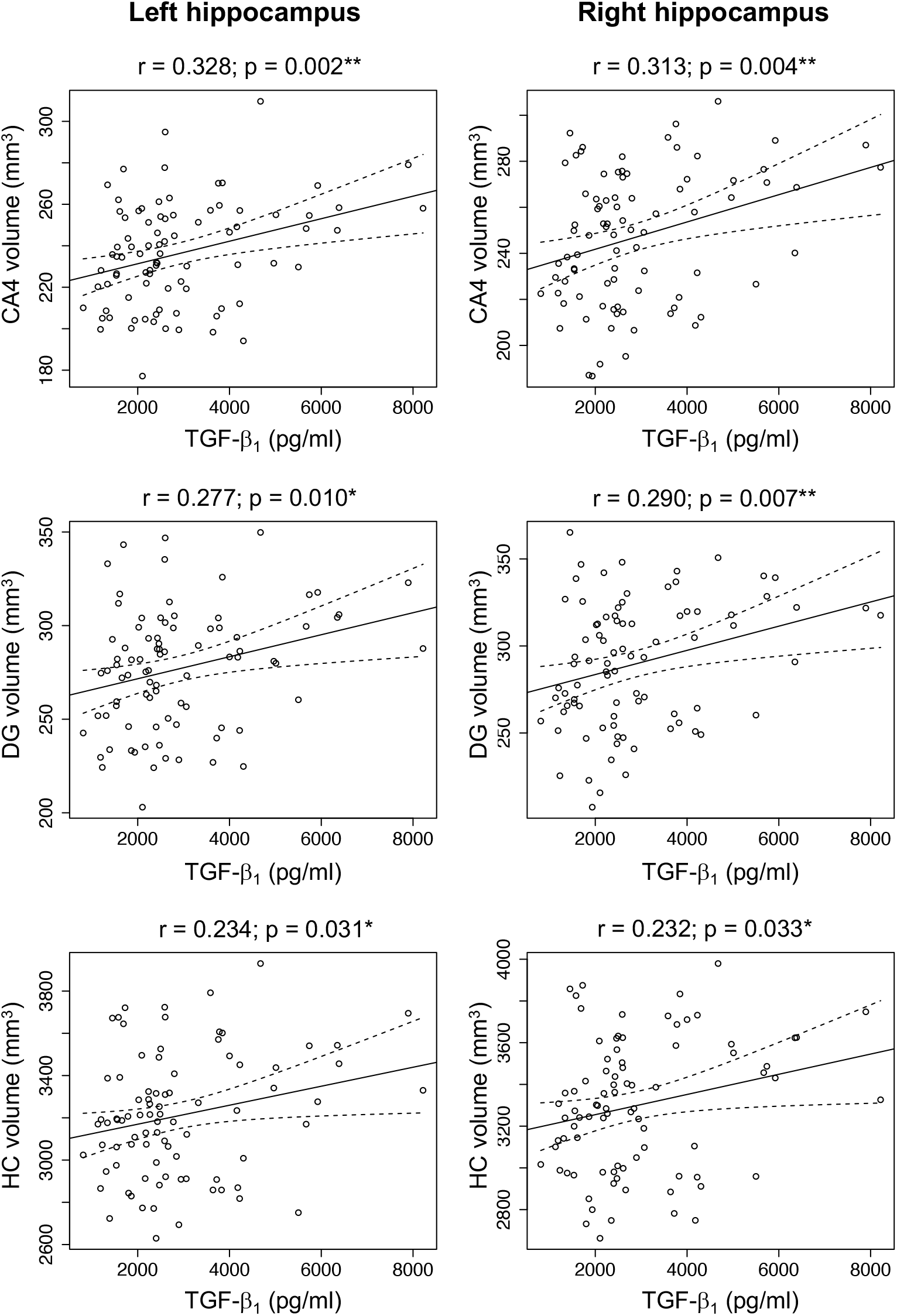
Correlation of TGF-β_1_ plasma concentrations with hippocampal subfield volumes. Pearson’s correlations are shown for the dentate gyrus (DG), CA4 and whole hippocampus, separated by hemisphere. * p < .05, two-tailed; ** p < .01, two-tailed.

### 3.3. Correlations of hippocampal subfield volumes with memory performance

To assess a potential relationship between the volumes of the hippocampus with memory performance in regions associated with immune markers, Pearson’s correlations between volumes and memory performance in the VLMT and WMS were computed for all regions showing a robust relationship (FDR-corrected p < .05) with at least one immune marker. As no robust associations were found in young adults, these analyses were restricted to the older participants in our cohort. We observed a positive correlation of the WMS Logical Memory scale with the volume of the right DG (p = .025, two-tailed) and, as a trend, also of left DG and the CA4 region bilaterally as well as the volume of the whole right hippocampus (all p < .05, one-tailed; see Figure 2). Moreover, significant positive correlations with memory performance in 24-hour delayed recall test of the WMS were observed for the volumes of the dentate gyrus bilaterally and the right CA4 region (all p < .05, two-tailed) and, as a trend, also for the left CA4 region (p = .042, two-tailed). On the other hand, for the VLMT, no significant correlations with the volumes of any of the hippocampal regions investigated were detected (all p > .096).

**Figure 2:**
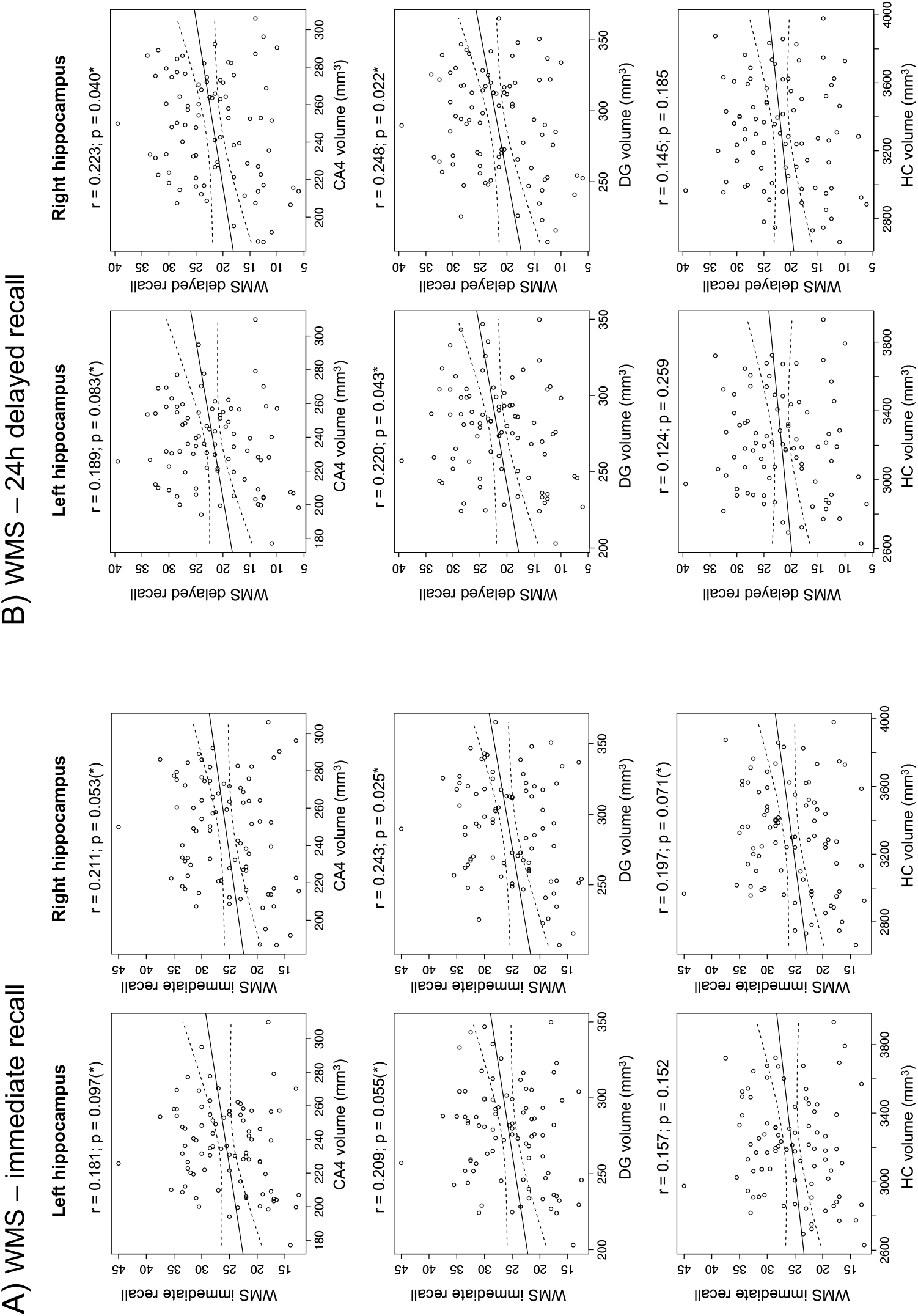
Correlation of hippocampal subfield volumes with memory performance (WMS Logical Memory). Pearson’s correlations are shown for the dentate gyrus (DG), CA4 and whole hippocampus, separated by hemisphere. A) WMS immediate recall. B) WMS 24 h delayed recall. * p < .05, two-tailed; (*) p < .05, one-tailed.

### 3.4. TGF-β plasma concentrations and memory performance

In both the learning and the 24 h delayed recall phases of the VLMT, regression analysis confirmed worse performance of older adults compared to young adults (linear regression, complete sample, effect of age; learning: standardized β = −0.70, confidence interval [CI] = [−0.83 −0.57], p < .001; delayed recall: standardized β = −0.66, confidence interval [CI] = [−0.80 −0.53], p < .001). Within the group of older adults, linear regression analysis revealed significant effects of age and gender on learning performance (age: standardized β = −0.34, CI = [−0.56 - 0.13], p = .002; gender: standardized β = 0.44, CI = [0.02 0.86], p = .038) and delayed recall (age: standardized β = −0.31, CI = [−0.53 −0.09], p = .006; gender: standardized β = 0.47, CI = [0.05 0.89], p = .028). We found, however, no effect of TGF-β1 plasma concentrations on learning or delayed recall performance (learning: standardized β = −0.00, CI = [-0.35 0.06], p = .157; delayed recall: standardized β = 0.00, CI = [-0.30 0.10], p = .328). There were also no effects of plasma levels of other cytokines or CRP nor of immune event history on either learning or delayed recall performance in the VLMT (all p > .217).

In the WMS, regression also confirmed worse performance in older versus young adults in both immediate and 24h delayed recall (complete sample, effect of age; immediate recall: standardized β = −0.46, confidence interval [CI] = [−0.63 −0.30], p < .001; delayed recall: standardized β = −0.48, confidence interval [CI] = [−0.64 −0.31], p < .001). Among older adults, we also found significant effects of age (immediate recall: standardized β = −0.38, CI = [−0.60 −0.17], p < .001; delayed recall: standardized β = −0.28, CI = [−0.51 −0.06], p = .014). As with the VLMT, however, no effect of TGF-β1 plasma concentrations on memory performance was found (immediate recall: standardized β = −0.00, CI = [-0.35 0.06], p = .157; delayed recall: standardized β = 0.00, CI = [-0.30 0.10], p = .328). There were also no significant effects of plasma levels of other cytokines or CRP or of immune event history on either immediate or delayed recall performance in the WMS, nor an effect of gender (all p > .165).

## 4. Discussion

Our results suggest that plasma concentrations of the anti-inflammatory cytokine TGF-β_1_ are associated with hippocampal CA4 and dentate gyrus volumes in older adults. The volumes of these hippocampal regions were in turn associated with immediate and particularly delayed recall performance in the WMS Logical Memory scale in the older participants.

### 4.1. A potentially protective role for TGF-β_1_ in the aging hippocampus

While young and older participants did not *per se* show age-related differences in TGF-β_1_ plasma concentrations (see Table 1), specifically in older adults, higher concentrations of TGF- β_1_ were associated with hippocampal structure and particularly the volumes of the hippocampal CA4 and DG subregions. Previous rodent studies and investigations in human neurodevelopmental disorders have pointed to a role for TGF-β signaling in hippocampal development and maintenance (Johnson et al. 2020; Stegeman et al. 2013; Oishi et al. 2016). In adult mice, TGF-β can promote hippocampal LTP, particularly the transition from early to late-phase LTP (Caraci et al. 2015; Nenov et al. 2019), as well as adult neurogenesis in the DG (Gradari et al. 2021). Given the age-dependence of the association between TGF-β_1_ plasma concentrations and hippocampal structure in the present study, one possibility could be that effects of impaired TGF-β signaling during development may be compensated for during young adulthood, but, during aging, the capacity for compensation decreases. On the other hand, it must be noted that TGF-β_1_ plasma concentrations were measured at or around the time of testing in our study, and we cannot make statements about long-term availability of TGF-β_1_.

Therefore, in our view, a more likely or perhaps additional explanation for the age-dependence of the observed association could be related to the prominent role of TGF-β_1_ signaling in CNS, but also systemic inflammation. It has been recognized for over two decades that microglia and astrocytes in particular play a central role as mediators between peripheral, low-grade inflammation and the cellular mechanisms of learning and memory (Yirmiya and Goshen 2011). They also express increased pro- and anti-inflammatory cytokines in aging and respond to peripheral inflammatory processes (Sierra et al. 2007). For example, microglia secrete the anti-inflammatory cytokine IL-10 in response to an inflammatory stimulus, which in turn induces astrocytes to produce TGF-β, resulting in an inhibition of the microglial inflammatory response (Norden et al. 2014). However, aging astrocytes frequently show a blunted response to IL-10, resulting in lower levels of TGF-β_1_ expression (Norden et al. 2016) and, consequently to a shift towards a pro-inflammatory state in microglia (Diniz et al. 2017), which has a negative impact on neural plasticity (Kempermann and Neumann 2003; Yirmiya and Goshen 2011). While this could in principle explain a relationship between higher TGF-β_1_ plasma levels and larger volumes of the hippocampal CA4 region and DG, an explanation primarily based on neurogenesis may be questionable, as recent investigations cast doubt on the robustness of findings on adult neurogenesis in humans (Sorrells et al. 2021). Nevertheless, the role of chronic inflammation in synaptic dysfunction and loss and ultimately neurodegeneration is a well-documented finding (Yirmiya and Goshen 2011; Daulatzai 2014), and increased TGF-β_1_ levels have been found in studies of pharmacological neuroprotection (Hosseini et al. 2018; Wiciński et al. 2018). Furthermore, TGF-β_1_ also promotes the outgrowth of dendrites (Battista et al. 2006; Luo et al. 2016; Mathieu et al. 2010).

One limitation of the present study is that we could measure TGF-β_1_ concentrations only in plasma but not, for example, in cerebrospinal fluid (CSF). There is surprisingly little available literature on the relationship between CSF and plasma concentrations of TGF-β_1_. Previous studies that included measures from both CSF and blood have often treated those measures separately and not reported the correlations between them (Iłzecka et al. 2002; Liu et al. 2022). One study in patients with Alzheimer’s disease explicitly mentioned an absence of such correlations (Motta et al. 2007), but in that study, peripheral blood levels of TGF-β_1_ were measured in serum and were thus most likely largely attributable to TGF-β_1_ released from degranulated platelets (Grainger et al. 2000; Iłzecka et al. 2002). In the present study, we used a protocol that avoids contamination from platelet-derived TGF-β_1_ (Reinhold et al. 1997), but a potential relationship with CSF TGF-β_1_ remains nevertheless elusive. Furthermore, it must be noted that TGF-β_1_ is not able to penetrate the intact blood-brain barrier (BBB) (Kastin et al. 2003). On the other hand, BBB integrity declines with age, even in healthy individuals. The hippocampus and in particular the CA1 region, but also the DG seem to be particularly affected by age-related BBB permeability, while the BBB remains comparably intact in neocortical areas (Montagne et al. 2015). Furthermore, increased BBB permeability in aging elicits increased activation of TGF-β signaling pathways and increased expression of TGF-β_1_ in the hippocampus (Senatorov et al. 2019). Additionally, TGF-β_1_ has been implicated in angiogenesis and BBB assembly (Diniz et al. 2017), and might increase reactively in response to BBB damage.

### 4.2. Hippocampal subregion structural integrity and memory performance in old age

While we observed no direct effect TGF-β_1_ plasma concentrations on memory performance, we found that the volumes of the hippocampal regions shown to be associated with TGF-β_1_ levels correlated with immediate and particularly delayed recall performance in the WMS Logical Memory scale in the older participants. Previous volumetric segmentation of hippocampal subregions and their association with mnestic abilities have revealed that the volumes of the input regions of the hippocampus (i.e., the DG and the CA4 and CA3 regions) are positively correlated with learning performance in well-established memory tests (Mueller et al. 2011; Aslaksen et al. 2018; Travis et al. 2014). On the other hand, the volumes of hippocampal output regions (i.e., CA1 and subiculum), are more related to delayed memory retrieval (Mueller et al. 2011; Aslaksen et al. 2018). This distinction is further supported by a functional MRI (fMRI) study on memory-related activations of hippocampal subregions (Eldridge et al. 2005). Nevertheless, caution is warranted, as other works have also yielded relationships between CA1 volume and learning performance as well as DG and CA4 volumes with performance in delayed retrieval (Aslaksen et al. 2018). Along the same line, a high-resolution fMRI study of successful encoding of information showed that encoding success correlated with activity in the CA1 region and subiculum, while activity in the DG, CA2 and CA3 was related to stimulus novelty (Maass et al. 2014). Evidence from rodent studies suggests that the volume of the CA4 region is associated with better spatial memory (Schwegler et al. 1990; Crusio and Schwegler 2005). To interpret these findings, one should keep in mind that “CA4” likely constitutes a misnomer, as, unlike CA1, CA3 and the small C2 region, CA4 mainly contains non-pyramidal cells and is functionally closely related to the adjacent DG (Amaral 1978; Amaral et al. 2007). As such, the region labeled “CA4” in the FreeSurfer-based segmentation of the hippocampus also contains the mossy fibers that originate from neurons of the DG, which was itself also associated with better memory performance in the older group, compatible with findings from previous studies (Zheng et al. 2018; Broadhouse et al. 2019; Kern et al. 2021; Wan et al. 2020). A larger volume in these regions might, for example, reflect both from more neurons in the DG due to better-preserved structural integrity and possibly neurogenesis as well as from a larger number of dendrites, possibly due to the role of TGF-β_1_ in promoting the outgrowth of dendrites (Battista et al. 2006; Luo et al. 2016; Mathieu et al. 2010).

### 4.3. Clinical implications and directions for future research

Our results suggest that plasma TGF-β_1_ levels are associated with a better-preserved hippocampal structure in older adults, which in turn is associated with greater episodic memory performance. Notably, our current data are based on physiological variability of TGF-β_1_ plasma levels, and potential effects of raising the concentrations to supra-physiological levels, for example, by pharmacological intervention, warrant further investigation. Increased TGF-β_1_ plasma levels have, for example, been found in patients with multiple sclerosis and are further increased by immunomodulatory treatment with interferon-β1b (Nicoletti et al. 1998), suggesting that the treatment might augment an already active endogenous anti-inflammatory mechanism. Little is known, however, about potential cognitive effects of elevated TGF-β_1_ levels. Notably, CSF TGF-β_1_ concentrations are increased in Alzheimer’s disease (Swardfager et al. 2010), but their relationship with cognitive performance depends on the disease stage, with both unaffected and severely affected individuals showing a positive correlation with cognitive performance (i.e., MMSE), whereas mildly to moderately affected patients exhibit a negative relationship (Motta et al. 2007). Future research should further elucidate the relationship between TGF-β_1_ levels and brain, particularly hippocampal, integrity in clinical populations to clarify, for example, its role in inflammatory processes related to neurodegeneration in Alzheimer’s disease (Brosseron et al. 2022).

Beyond TGF-β, our results provide further evidence for a relationship between peripheral markers of inflammation and brain structure, particularly hippocampal structure, as well as neurocognitive functioning (Baune et al. 2012; Marsland et al. 2008; Yirmiya and Goshen 2011; Brosseron et al. 2023). Future studies should expand those works, for example, by employing functional MRI and derived biomarkers (Soch et al. 2021a; Richter et al. 2023). Another important direction for future research concerns gender differences in inflammaging. In the present study, we observed higher CRP levels in young, but not older, women compared to men, which have been reported previously by other groups and partly attributed to gender differences in body fat or estrogen levels (Khera et al. 2005; Khera et al. 2009; Lakoski et al. 2006). As we found no gender differences in older adults, future studies should explore the possibility that, for example, hormonal differences and their lifespan-associated changes might contribute to immunosenescence and associated neurocognitive alterations.

### 4.5. Conclusions

Our results suggest that TGF-β1 plasma levels are associated with better structural preservation of the hippocampus in older adults, which is predictive for better episodic memory performance. The present data highlight the role of anti-inflammatory mechanisms as potential protective factors in neurocognitive aging.

## 5. Notes

## 5.1. Acknowledgments

The authors would like to thank Hannah Feldhoff for help with neuropsychological data collection and Kerstin Möhring, Ilona Wiedenhöft, and Claus Tempelmann for expert technical assistance with MRI data acquisition.

This study was supported by the State of Saxony-Anhalt and the European Union (Research Alliance “Autonomy in Old Age” to B.H.S., D.R., B.S., and I.R.D.) and by the Deutsche Forschungsgemeinschaft (DFG, German Research Foundation) - 425899996/CRC1436 to B.H.S. and C.I.S.; SFB854, to B.S and IRD; 362321501/RTG 2413 SynAGE to C.I.S.). B.S also receives additional funding by grants from the state of Saxony-Anhalt (SI-2 and SI-3). The funding agencies had no role in the design or analysis of the study. The authors have no conflict of interest, financial or otherwise, to declare.

## 5.2. Data Availability Statement

Access to de-identified raw data and R scripts used for data analysis will be provided by the authors upon reasonable request.

## References

Abe K, Chu PJ, Ishihara A, Saito H (1996) Transforming growth factor-beta 1 promotes re-elongation of injured axons of cultured rat hippocampal neurons. Brain Res 723 (1-2):206–209. doi: 10.1016/0006-8993(96)00253-3

Ackenheil M, Stotz G, Dietz-Bauer R, Vossen A, Dietz R, Vossen-Wellmann A, Vossen JA (1999) Mini International Neuropsychiatric Interview. German Version 5.0.0, DSM-IV. Psychiatrische Universitätsklinik München, München

Acosta JC, Banito A, Wuestefeld T, Georgilis A, Janich P, Morton JP, Athineos D, Kang TW, Lasitschka F, Andrulis M, Pascual G, Morris KJ, Khan S, Jin H, Dharmalingam G, Snijders AP, Carroll T, Capper D, Pritchard C, Inman GJ, Longerich T, Sansom OJ, Benitah SA, Zender L, Gil J (2013) A complex secretory program orchestrated by the inflammasome controls paracrine senescence. Nat Cell Biol 15 (8):978–990. doi: 10.1038/ncb2784

Akbaraly TN, Hamer M, Ferrie JE, Lowe G, Batty GD, Hagger-Johnson G, Singh-Manoux A, Shipley MJ, Kivimäki M (2013) Chronic inflammation as a determinant of future aging phenotypes. Cmaj 185 (16):E763–770. doi: 10.1503/cmaj.122072

Amaral DG (1978) A Golgi study of cell types in the hilar region of the hippocampus in the rat. J Comp Neurol 182 (4 Pt 2):851–914. doi: 10.1002/cne.901820508

Amaral DG, Scharfman HE, Lavenex P (2007) The dentate gyrus: fundamental neuroanatomical organization (dentate gyrus for dummies). Prog Brain Res 163:3–22. doi: 10.1016/s0079-6123(07)63001-5

Aslaksen PM, Bystad MK, Ørbo MC, Vangberg TR (2018) The relation of hippocampal subfield volumes to verbal episodic memory measured by the California Verbal Learning Test II in healthy adults. Behav Brain Res 351:131–137. doi: 10.1016/j.bbr.2018.06.008

Basisty N, Kale A, Jeon OH, Kuehnemann C, Payne T, Rao C, Holtz A, Shah S, Sharma V, Ferrucci L, Campisi J, Schilling B (2020) A proteomic atlas of senescence-associated secretomes for aging biomarker development. PLoS Biol 18 (1):e3000599. doi: 10.1371/journal.pbio.3000599

Battista D, Ferrari CC, Gage FH, Pitossi FJ (2006) Neurogenic niche modulation by activated microglia: transforming growth factor beta increases neurogenesis in the adult dentate gyrus. Eur J Neurosci 23 (1):83–93. doi: 10.1111/j.1460-9568.2005.04539.x

Baune BT, Konrad C, Grotegerd D, Suslow T, Birosova E, Ohrmann P, Bauer J, Arolt V, Heindel W, Domschke K, Schöning S, Rauch AV, Uhlmann C, Kugel H, Dannlowski U (2012) Interleukin-6 gene (IL-6): a possible role in brain morphology in the healthy adult brain. J Neuroinflammation 9:125. doi: 10.1186/1742-2094-9-125

Bond AM, Ming GL, Song H (2021) Ontogeny of adult neural stem cells in the mammalian brain. Curr Top Dev Biol 142:67–98. doi: 10.1016/bs.ctdb.2020.11.002

Brionne TC, Tesseur I, Masliah E, Wyss-Coray T (2003) Loss of TGF-beta 1 leads to increased neuronal cell death and microgliosis in mouse brain. Neuron 40 (6):1133–1145. doi: 10.1016/s0896-6273(03)00766-9

Broadhouse KM, Mowszowski L, Duffy S, Leung I, Cross N, Valenzuela MJ, Naismith SL (2019) Memory Performance Correlates of Hippocampal Subfield Volume in Mild Cognitive Impairment Subtype. Front Behav Neurosci 13:259. doi: 10.3389/fnbeh.2019.00259

Brosseron F, Maass A, Kleineidam L, Ravichandran KA, González PG, McManus RM, Ising C, Santarelli F, Kolbe CC, Häsler LM, Wolfsgruber S, Marquié M, Boada M, Orellana A, de Rojas I, Röske S, Peters O, Cosma NC, Cetindag A, Wang X, Priller J, Spruth EJ, Altenstein S, Schneider A, Fliessbach K, Wiltfang J, Schott BH, Bürger K, Janowitz D, Dichgans M, Perneczky R, Rauchmann BS, Teipel S, Kilimann I, Goerss D, Laske C, Munk MH, Düzel E, Yakupov R, Dobisch L, Metzger CD, Glanz W, Ewers M, Dechent P, Haynes JD, Scheffler K, Roy N, Rostamzadeh A, Teunissen CE, Marchant NL, Spottke A, Jucker M, Latz E, Wagner M, Mengel D, Synofzik M, Jessen F, Ramirez A, Ruiz A, Heneka MT (2022) Soluble TAM receptors sAXL and sTyro3 predict structural and functional protection in Alzheimer’s disease. Neuron 110 (6):1009–1022.e1004. doi: 10.1016/j.neuron.2021.12.016

Brosseron F, Maass A, Kleineidam L, Ravichandran KA, Kolbe CC, Wolfsgruber S, Santarelli F, Häsler LM, McManus R, Ising C, Röske S, Peters O, Cosma NC, Schneider LS, Wang X, Priller J, Spruth EJ, Altenstein S, Schneider A, Fliessbach K, Wiltfang J, Schott BH, Buerger K, Janowitz D, Dichgans M, Perneczky R, Rauchmann BS, Teipel S, Kilimann I, Görß D, Laske C, Munk MH, Düzel E, Yakupow R, Dobisch L, Metzger CD, Glanz W, Ewers M, Dechent P, Haynes JD, Scheffler K, Roy N, Rostamzadeh A, Spottke A, Ramirez A, Mengel D, Synofzik M, Jucker M, Latz E, Jessen F, Wagner M, Heneka MT (2023) Serum IL-6, sAXL, and YKL-40 as systemic correlates of reduced brain structure and function in Alzheimer’s disease: results from the DELCODE study. Alzheimers Res Ther 15 (1):13. doi: 10.1186/s13195-022-01118-0

Caraci F, Gulisano W, Guida CA, Impellizzeri AA, Drago F, Puzzo D, Palmeri A (2015) A key role for TGF-β1 in hippocampal synaptic plasticity and memory. Sci Rep 5:11252. doi: 10.1038/srep11252

Chen JH, Ke KF, Lu JH, Qiu YH, Peng YP (2015) Protection of TGF-β1 against neuroinflammation and neurodegeneration in Aβ1-42-induced Alzheimer’s disease model rats. PLoS One 10 (2):e0116549. doi: 10.1371/journal.pone.0116549

Crusio WE, Schwegler H (2005) Learning spatial orientation tasks in the radial-maze and structural variation in the hippocampus in inbred mice. Behav Brain Funct 1 (1):3. doi: 10.1186/1744-9081-1-3

Daulatzai MA (2014) Role of stress, depression, and aging in cognitive decline and Alzheimer’s disease. Curr Top Behav Neurosci 18:265–296. doi: 10.1007/7854_2014_350

De Martinis M, Franceschi C, Monti D, Ginaldi L (2005) Inflamm-ageing and lifelong antigenic load as major determinants of ageing rate and longevity. FEBS Lett 579 (10):2035–2039. doi: 10.1016/j.febslet.2005.02.055

Diniz LP, Tortelli V, Matias I, Morgado J, Bérgamo Araujo AP, Melo HM, Seixas da Silva GS, Alves-Leon SV, de Souza JM, Ferreira ST, De Felice FG, Gomes FCA (2017) Astrocyte Transforming Growth Factor Beta 1 Protects Synapses against Aβ Oligomers in Alzheimer’s Disease Model. J Neurosci 37 (28):6797–6809. doi: 10.1523/jneurosci.3351-16.2017

Dobaczewski M, Chen W, Frangogiannis NG (2011) Transforming growth factor (TGF)-β signaling in cardiac remodeling. J Mol Cell Cardiol 51 (4):600–606. doi: 10.1016/j.yjmcc.2010.10.033

Dounavi ME, Mak E, Wells K, Ritchie K, Ritchie CW, Su L, JT OB (2020) Volumetric alterations in the hippocampal subfields of subjects at increased risk of dementia. Neurobiol Aging 91:36–44. doi: 10.1016/j.neurobiolaging.2020.03.006

Eldridge LL, Engel SA, Zeineh MM, Bookheimer SY, Knowlton BJ (2005) A dissociation of encoding and retrieval processes in the human hippocampus. J Neurosci 25 (13):3280–3286. doi: 10.1523/jneurosci.3420-04.2005

Fischl B (2012) FreeSurfer. Neuroimage 62 (2):774–781. doi: 10.1016/j.neuroimage.2012.01.021

Folstein MF, Folstein SE, McHugh PR (1975) “Mini-mental state”. A practical method for grading the cognitive state of patients for the clinician. J Psychiatr Res 12 (3):189–198. doi: 10.1016/0022-3956(75)90026-6

Franceschi C (2007) Inflammaging as a major characteristic of old people: can it be prevented or cured? Nutr Rev 65 (12 Pt 2):S173–176. doi: 10.1111/j.1753-4887.2007.tb00358.x

Franceschi C, Bonafè M, Valensin S, Olivieri F, De Luca M, Ottaviani E, De Benedictis G (2000) Inflamm-aging. An evolutionary perspective on immunosenescence. Ann N Y Acad Sci 908:244–254. doi: 10.1111/j.1749-6632.2000.tb06651.x

Fülöp T, Herbein G, Cossarizza A, Witkowski JM, Frost E, Dupuis G, Pawelec G, Larbi A (2017) Cellular Senescence, Immunosenescence and HIV. Interdiscip Top Gerontol Geriatr 42:28–46. doi: 10.1159/000448542

Gorbach T, Pudas S, Lundquist A, Orädd G, Josefsson M, Salami A, de Luna X, Nyberg L (2017) Longitudinal association between hippocampus atrophy and episodic-memory decline. Neurobiol Aging 51:167–176. doi: 10.1016/j.neurobiolaging.2016.12.002

Gradari S, Herrera A, Tezanos P, Fontán-Lozano Á, Pons S, Trejo JL (2021) The Role of Smad2 in Adult Neuroplasticity as Seen through Hippocampal-Dependent Spatial Learning/Memory and Neurogenesis. J Neurosci 41 (32):6836–6849. doi: 10.1523/jneurosci.2619-20.2021

Grainger DJ, Mosedale DE, Metcalfe JC (2000) TGF-beta in blood: a complex problem. Cytokine Growth Factor Rev 11 (1-2):133–145. doi: 10.1016/s1359-6101(99)00037-4

Harder T, Guttek K, Philipsen L, Simeoni L, Schraven B, Reinhold D (2014) Selective targeting of transforming growth factor-beta1 into TCR/CD28 signalling plasma membrane domains silences T cell activation. Cell Commun Signal 12:74. doi: 10.1186/s12964-014-0074-6

Härting C, Markowitsch H-J, Neufeld H, Calabrese P, Deisinger K, Kessler J (2000) Wechsler Memory Scale – Revised Edition, German Edition. Manual. Huber, Bern

Helmstaedter C (2001) VLMT: Verbaler Lern-und Merkfähigkeitstest. . Beltz Test, Göttingen:

Hosseini SM, Gholami Pourbadie H, Sayyah M, Zibaii MI, Naderi N (2018) Neuroprotective effect of monophosphoryl lipid A, a detoxified lipid A derivative, in photothrombotic model of unilateral selective hippocampal ischemia in rat. Behav Brain Res 347:26–36. doi: 10.1016/j.bbr.2018.02.045

Iglesias JE, Augustinack JC, Nguyen K, Player CM, Player A, Wright M, Roy N, Frosch MP, McKee AC, Wald LL, Fischl B, Van Leemput K (2015) A computational atlas of the hippocampal formation using ex vivo, ultra-high resolution MRI: Application to adaptive segmentation of in vivo MRI. Neuroimage 115:117–137. doi: 10.1016/j.neuroimage.2015.04.042

Iłzecka J, Stelmasiak Z, Dobosz B (2002) Transforming growth factor-Beta 1 (tgf-Beta 1) in patients with amyotrophic lateral sclerosis. Cytokine 20 (5):239–243. doi: 10.1006/cyto.2002.2005

Johnson BV, Kumar R, Oishi S, Alexander S, Kasherman M, Vega MS, Ivancevic A, Gardner A, Domingo D, Corbett M, Parnell E, Yoon S, Oh T, Lines M, Lefroy H, Kini U, Van Allen M, Grønborg S, Mercier S, Küry S, Bézieau S, Pasquier L, Raynaud M, Afenjar A, Billette de Villemeur T, Keren B, Désir J, Van Maldergem L, Marangoni M, Dikow N, Koolen DA, VanHasselt PM, Weiss M, Zwijnenburg P, Sa J, Reis CF, López-Otín C, Santiago-Fernández O, Fernández-Jaén A, Rauch A, Steindl K, Joset P, Goldstein A, Madan-Khetarpal S, Infante E, Zackai E, McDougall C, Narayanan V, Ramsey K, Mercimek-Andrews S, Pena L, Shashi V, Schoch K, Sullivan JA, Pinto EVF, Pichurin PN, Ewing SA, Barnett SS, Klee EW, Perry MS, Koenig MK, Keegan CE, Schuette JL, Asher S, Perilla-Young Y, Smith LD, Rosenfeld JA, Bhoj E, Kaplan P, Li D, Oegema R, van Binsbergen E, van der Zwaag B, Smeland MF, Cutcutache I, Page M, Armstrong M, Lin AE, Steeves MA, Hollander ND, Hoffer MJV, Reijnders MRF, Demirdas S, Koboldt DC, Bartholomew D, Mosher TM, Hickey SE, Shieh C, Sanchez-Lara PA, Graham JM, Jr., Tezcan K, Schaefer GB, Danylchuk NR, Asamoah A, Jackson KE, Yachelevich N, Au M, Pérez-Jurado LA, Kleefstra T, Penzes P, Wood SA, Burne T, Pierson TM, Piper M, Gécz J, Jolly LA (2020) Partial Loss of USP9X Function Leads to a Male Neurodevelopmental and Behavioral Disorder Converging on Transforming Growth Factor β Signaling. Biol Psychiatry 87 (2):100–112. doi: 10.1016/j.biopsych.2019.05.028

Kastin AJ, Akerstrom V, Pan W (2003) Circulating TGF-beta1 does not cross the intact blood-brain barrier. J Mol Neurosci 21 (1):43–48. doi: 10.1385/jmn:21:1:43

Kempermann G, Neumann H (2003) Neuroscience. Microglia: the enemy within? Science 302 (5651):1689–1690. doi: 10.1126/science.1092864

Kern KL, Storer TW, Schon K (2021) Cardiorespiratory fitness, hippocampal subfield volumes, and mnemonic discrimination task performance in aging. Hum Brain Mapp 42 (4):871–892. doi: 10.1002/hbm.25259

Khera A, McGuire DK, Murphy SA, Stanek HG, Das SR, Vongpatanasin W, Wians FH, Jr., Grundy SM, de Lemos JA (2005) Race and gender differences in C-reactive protein levels. J Am Coll Cardiol 46 (3):464–469. doi: 10.1016/j.jacc.2005.04.051

Khera A, Vega GL, Das SR, Ayers C, McGuire DK, Grundy SM, de Lemos JA (2009) Sex differences in the relationship between C-reactive protein and body fat. J Clin Endocrinol Metab 94 (9):3251–3258. doi: 10.1210/jc.2008-2406

Kirchner K, Garvert L, Wittfeld K, Ameling S, Bülow R, Meyer Zu Schwabedissen H, Nauck M, Völzke H, Grabe HJ, Van der Auwera S (2023) Deciphering the Effect of Different Genetic Variants on Hippocampal Subfield Volumes in the General Population. Int J Mol Sci 24 (2). doi: 10.3390/ijms24021120

Lakoski SG, Cushman M, Criqui M, Rundek T, Blumenthal RS, D’Agostino RB, Jr., Herrington DM (2006) Gender and C-reactive protein: data from the Multiethnic Study of Atherosclerosis (MESA) cohort. Am Heart J 152 (3):593–598. doi: 10.1016/j.ahj.2006.02.015

Lehrl S (2005) Mehrfachwahl-Wortschatz-Intelligenztest MWT-B 5th ed. edn. Spitta,

Liu YC, Hsiao HT, Wang JC, Wen TC, Chen SL (2022) TGF-β1 in plasma and cerebrospinal fluid can be used as a biological indicator of chronic pain in patients with osteoarthritis. PLoS One 17 (1):e0262074. doi: 10.1371/journal.pone.0262074

Lüdecke D (2022) sjPlot: Data Visualization for Statistics in Social Science. R package version 2.8.12

Luo SX, Timbang L, Kim JI, Shang Y, Sandoval K, Tang AA, Whistler JL, Ding JB, Huang EJ (2016) TGF-β Signaling in Dopaminergic Neurons Regulates Dendritic Growth, Excitatory-Inhibitory Synaptic Balance, and Reversal Learning. Cell Rep 17 (12):3233–3245. doi: 10.1016/j.celrep.2016.11.068

Maass A, Schütze H, Speck O, Yonelinas A, Tempelmann C, Heinze HJ, Berron D, Cardenas-Blanco A, Brodersen KH, Stephan KE, Düzel E (2014) Laminar activity in the hippocampus and entorhinal cortex related to novelty and episodic encoding. Nat Commun 5:5547. doi: 10.1038/ncomms6547

Marsland AL, Gianaros PJ, Abramowitch SM, Manuck SB, Hariri AR (2008) Interleukin-6 covaries inversely with hippocampal grey matter volume in middle-aged adults. Biol Psychiatry 64 (6):484–490. doi: 10.1016/j.biopsych.2008.04.016

Martínez-Canabal A (2015) Potential neuroprotective role of transforming growth factor β1 (TGFβ1) in the brain. Int J Neurosci 125 (1):1–9. doi: 10.3109/00207454.2014.903947

Mathieu P, Piantanida AP, Pitossi F (2010) Chronic expression of transforming growth factor-beta enhances adult neurogenesis. Neuroimmunomodulation 17 (3):200–201. doi: 10.1159/000258723

Mikheeva IB, Malkov AE, Pavlik LL, Arkhipov VI, Levin SG (2019) Effect of TGF-beta1 on long-term synaptic plasticity and distribution of AMPA receptors in the CA1 field of the hippocampus. Neurosci Lett 704:95–99. doi: 10.1016/j.neulet.2019.04.005

Montagne A, Barnes SR, Sweeney MD, Halliday MR, Sagare AP, Zhao Z, Toga AW, Jacobs RE, Liu CY, Amezcua L, Harrington MG, Chui HC, Law M, Zlokovic BV (2015) Blood-brain barrier breakdown in the aging human hippocampus. Neuron 85 (2):296–302. doi: 10.1016/j.neuron.2014.12.032

Motta M, Imbesi R, Di Rosa M, Stivala F, Malaguarnera L (2007) Altered plasma cytokine levels in Alzheimer’s disease: correlation with the disease progression. Immunol Lett 114 (1):46–51. doi: 10.1016/j.imlet.2007.09.002

Mueller SG, Chao LL, Berman B, Weiner MW (2011) Evidence for functional specialization of hippocampal subfields detected by MR subfield volumetry on high resolution images at 4 T. Neuroimage 56 (3):851–857. doi: 10.1016/j.neuroimage.2011.03.028

Nelson G, Wordsworth J, Wang C, Jurk D, Lawless C, Martin-Ruiz C, von Zglinicki T (2012) A senescent cell bystander effect: senescence-induced senescence. Aging Cell 11 (2):345–349. doi: 10.1111/j.1474-9726.2012.00795.x

Nenov MN, Malkov AE, Konakov MV, Levin SG (2019) Interleukin-10 and transforming growth factor-β1 facilitate long-term potentiation in CA1 region of hippocampus. Biochem Biophys Res Commun 518 (3):486–491. doi: 10.1016/j.bbrc.2019.08.072

Nicoletti F, Di Marco R, Patti F, Reggio E, Nicoletti A, Zaccone P, Stivala F, Meroni PL, Reggio A (1998) Blood levels of transforming growth factor-beta 1 (TGF-beta1) are elevated in both relapsing remitting and chronic progressive multiple sclerosis (MS) patients and are further augmented by treatment with interferon-beta 1b (IFN-beta1b). Clin Exp Immunol 113 (1):96–99. doi: 10.1046/j.1365-2249.1998.00604.x

Norden DM, Fenn AM, Dugan A, Godbout JP (2014) TGFβ produced by IL-10 redirected astrocytes attenuates microglial activation. Glia 62 (6):881–895. doi: 10.1002/glia.22647

Norden DM, Trojanowski PJ, Walker FR, Godbout JP (2016) Insensitivity of astrocytes to interleukin 10 signaling following peripheral immune challenge results in prolonged microglial activation in the aged brain. Neurobiol Aging 44:22–41. doi: 10.1016/j.neurobiolaging.2016.04.014

Nyberg L (2017) Functional brain imaging of episodic memory decline in ageing. J Intern Med 281 (1):65–74. doi: 10.1111/joim.12533

Nyberg L, Pudas S (2019) Successful Memory Aging. Annu Rev Psychol 70:219–243. doi: 10.1146/annurev-psych-010418-103052

Oishi S, Premarathne S, Harvey TJ, Iyer S, Dixon C, Alexander S, Burne TH, Wood SA, Piper M (2016) Usp9x-deficiency disrupts the morphological development of the postnatal hippocampal dentate gyrus. Sci Rep 6:25783. doi: 10.1038/srep25783

Preller V, Gerber A, Wrenger S, Togni M, Marguet D, Tadje J, Lendeckel U, Röcken C, Faust J, Neubert K, Schraven B, Martin R, Ansorge S, Brocke S, Reinhold D (2007) TGF-beta1-mediated control of central nervous system inflammation and autoimmunity through the inhibitory receptor CD26. J Immunol 178 (7):4632–4640. doi: 10.4049/jimmunol.178.7.4632

Quattrini G, Pievani M, Jovicich J, Aiello M, Bargalló N, Barkhof F, Bartres-Faz D, Beltramello A, Pizzini FB, Blin O, Bordet R, Caulo M, Constantinides M, Didic M, Drevelegas A, Ferretti A, Fiedler U, Floridi P, Gros-Dagnac H, Hensch T, Hoffmann KT, Kuijer JP, Lopes R, Marra C, Müller BW, Nobili F, Parnetti L, Payoux P, Picco A, Ranjeva JP, Roccatagliata L, Rossini PM, Salvatore M, Schonknecht P, Schott BH, Sein J, Soricelli A, Tarducci R, Tsolaki M, Visser PJ, Wiltfang J, Richardson JC, Frisoni GB, Marizzoni M (2020) Amygdalar nuclei and hippocampal subfields on MRI: Test-retest reliability of automated volumetry across different MRI sites and vendors. Neuroimage 218:116932. doi: 10.1016/j.neuroimage.2020.116932

Rattan SI (2014) Aging is not a disease: implications for intervention. Aging Dis 5 (3):196–202. doi: 10.14336/ad.2014.0500196

Reinhold D, Bank U, Bühling F, Junker U, Kekow J, Schleicher E, Ansorge S (1997) A detailed protocol for the measurement of TGF-beta1 in human blood samples. J Immunol Methods 209 (2):203–206. doi: 10.1016/s0022-1759(97)00160-9

Revelle W (2022) psych: Procedures for Psychological, Psychometric, and Personality Research. R package version 2.2.9 Northwestern University, Evanston, Illinois.

Richter A, Soch J, Kizilirmak JM, Fischer L, Schütze H, Assmann A, Behnisch G, Feldhoff H, Knopf L, Raschick M, Schult A, Seidenbecher CI, Yakupov R, Düzel E, Schott BH (2023) Single-value scores of memory-related brain activity reflect dissociable neuropsychological and anatomical signatures of neurocognitive aging. Hum Brain Mapp. doi: 10.1002/hbm.26281

Schwegler H, Crusio WE, Brust I (1990) Hippocampal mossy fibers and radial-maze learning in the mouse: a correlation with spatial working memory but not with non-spatial reference memory. Neuroscience 34 (2):293–298. doi: 10.1016/0306-4522(90)90139-u

Senatorov VV, Jr., Friedman AR, Milikovsky DZ, Ofer J, Saar-Ashkenazy R, Charbash A, Jahan N, Chin G, Mihaly E, Lin JM, Ramsay HJ, Moghbel A, Preininger MK, Eddings CR, Harrison HV, Patel R, Shen Y, Ghanim H, Sheng H, Veksler R, Sudmant PH, Becker A, Hart B, Rogawski MA, Dillin A, Friedman A, Kaufer D (2019) Blood-brain barrier dysfunction in aging induces hyperactivation of TGFβ signaling and chronic yet reversible neural dysfunction. Sci Transl Med 11 (521). doi: 10.1126/scitranslmed.aaw8283

Sheehan DV, Lecrubier Y, Sheehan KH, Amorim P, Janavs J, Weiller E, Hergueta T, Baker R, Dunbar GC (1998) The Mini-International Neuropsychiatric Interview (M.I.N.I.): the development and validation of a structured diagnostic psychiatric interview for DSM-IV and ICD-10. J Clin Psychiatry 59 Suppl 20:22–-33;quiz 34-57

Shen WX, Chen JH, Lu JH, Peng YP, Qiu YH (2014) TGF-β1 protection against Aβ1-42-induced neuroinflammation and neurodegeneration in rats. Int J Mol Sci 15 (12):22092–22108. doi: 10.3390/ijms151222092

Sierra A, Gottfried-Blackmore AC, McEwen BS, Bulloch K (2007) Microglia derived from aging mice exhibit an altered inflammatory profile. Glia 55 (4):412–424. doi: 10.1002/glia.20468

Soch J, Richter A, Schütze H, Kizilirmak JM, Assmann A, Behnisch G, Feldhoff H, Fischer L, Heil J, Knopf L, Merkel C, Raschick M, Schietke CJ, Schult A, Seidenbecher CI, Yakupov R, Ziegler G, Wiltfang J, Düzel E, Schott BH (2021a) A comprehensive score reflecting memory-related fMRI activations and deactivations as potential biomarker for neurocognitive aging. Hum Brain Mapp 42 (14):4478–4496. doi: 10.1002/hbm.25559

Soch J, Richter A, Schütze H, Kizilirmak JM, Assmann A, Knopf L, Raschick M, Schult A, Maass A, Ziegler G, Richardson-Klavehn A, Düzel E, Schott BH (2021b) Bayesian model selection favors parametric over categorical fMRI subsequent memory models in young and older adults. Neuroimage 230:117820. doi: 10.1016/j.neuroimage.2021.117820

Sorrells SF, Paredes MF, Zhang Z, Kang G, Pastor-Alonso O, Biagiotti S, Page CE, Sandoval K, Knox A, Connolly A, Huang EJ, Garcia-Verdugo JM, Oldham MC, Yang Z, Alvarez-Buylla A (2021) Positive Controls in Adults and Children Support That Very Few, If Any, New Neurons Are Born in the Adult Human Hippocampus. J Neurosci 41 (12):2554–2565. doi: 10.1523/jneurosci.0676-20.2020

Stegeman S, Jolly LA, Premarathne S, Gecz J, Richards LJ, Mackay-Sim A, Wood SA (2013) Loss of Usp9x disrupts cortical architecture, hippocampal development and TGFβ-mediated axonogenesis. PLoS One 8 (7):e68287. doi: 10.1371/journal.pone.0068287

Swardfager W, Lanctôt K, Rothenburg L, Wong A, Cappell J, Herrmann N (2010) A meta-analysis of cytokines in Alzheimer’s disease. Biol Psychiatry 68 (10):930–941. doi: 10.1016/j.biopsych.2010.06.012

Tan A, Ma W, Vira A, Marwha D, Eliot L (2016) The human hippocampus is not sexually-dimorphic: Meta-analysis of structural MRI volumes. Neuroimage 124 (Pt A):350–366. doi: 10.1016/j.neuroimage.2015.08.050

Tracy RP (2003) Emerging relationships of inflammation, cardiovascular disease and chronic diseases of aging. Int J Obes Relat Metab Disord 27 Suppl 3:S29–34. doi: 10.1038/sj.ijo.0802497

Travis SG, Huang Y, Fujiwara E, Radomski A, Olsen F, Carter R, Seres P, Malykhin NV (2014) High field structural MRI reveals specific episodic memory correlates in the subfields of the hippocampus. Neuropsychologia 53:233–245. doi: 10.1016/j.neuropsychologia.2013.11.016

Wan M, Ye Y, Lin H, Xu Y, Liang S, Xia R, He J, Qiu P, Huang C, Tao J, Chen L, Zheng G (2020) Deviations in Hippocampal Subregion in Older Adults With Cognitive Frailty. Front Aging Neurosci 12:615852. doi: 10.3389/fnagi.2020.615852

Wiciński M, Wódkiewicz E, Słupski M, Walczak M, Socha M, Malinowski B, Pawlak-Osińska K (2018) Neuroprotective Activity of Sitagliptin via Reduction of Neuroinflammation beyond the Incretin Effect: Focus on Alzheimer’s Disease. Biomed Res Int 2018:6091014. doi: 10.1155/2018/6091014

World Medical Association Declaration of Helsinki: ethical principles for medical research involving human subjects (2013). Jama 310 (20):2191–2194. doi: 10.1001/jama.2013.281053

Yirmiya R, Goshen I (2011) Immune modulation of learning, memory, neural plasticity and neurogenesis. Brain Behav Immun 25 (2):181–213. doi: 10.1016/j.bbi.2010.10.015

Zheng F, Cui D, Zhang L, Zhang S, Zhao Y, Liu X, Liu C, Li Z, Zhang D, Shi L, Liu Z, Hou K, Lu W, Yin T, Qiu J (2018) The Volume of Hippocampal Subfields in Relation to Decline of Memory Recall Across the Adult Lifespan. Front Aging Neurosci 10:320. doi: 10.3389/fnagi.2018.00320

